# Unsupervised classification of SARS-CoV-2 genomic sequences uncovers hidden genetic diversity and suggests an efficient strategy for genomic surveillance

**DOI:** 10.1101/2021.06.23.449558

**Authors:** Matteo Chiara, David S. Horner, Erika Ferrandi, Carmela Gissi, Graziano Pesole

**Affiliations:** Department of Biosciences, University of Milan, Italy; Institute of Biomembranes, Bioenergetics and Molecular Biotechnologies, Consiglio Nazionale delle Ricerche, Bari, Italy; Department of Biosciences, Biotechnology and Biopharmaceutics, University of Bari “A. Moro”, Italy

**Keywords:** SARS-CoV-2, genomic surveillance, emerging mutations, classification, comparative genomics

## Abstract

Accurate and timely monitoring of emerging genomic diversity is crucial for limiting the spread of potentially more transmissible/pathogenic strains of SARS-CoV-2. At the time of writing, over 1.8M distinct viral genome sequences have been made publicly available, and a sophisticated nomenclature system based on phylogenetic evidence and expert manual curation has allowed the relatively rapid classification of emerging lineages of potential concern.

Here, we propose a complementary approach that integrates fine-grained spatiotemporal estimates of allele frequency with unsupervised clustering of viral haplotypes, and demonstrate that multiple highly frequent genetic variants, arising within large and/or rapidly expanding SARS-CoV-2 lineages, have highly biased geographic distributions and are not adequately captured by current SARS-CoV-2 nomenclature standards.

Our results advocate a partial revision of current methods used to track SARS-CoV-2 genomic diversity and highlight the importance of the application of strategies based on the systematic analysis and integration of regional data. Here we provide a complementary, completely automated and reproducible framework for the mapping of genetic diversity in time and across different geographic regions, and for the prioritization of virus variants of potential concern. We believe that the approach outlined in this study will contribute to relevant advances to current genomic surveillance methods.

## Main

The timely and accurate identification of emerging variants of a pathogen is one of the most important and ambitious goals of genomic surveillance strategies and represents the first line of defense for limiting the spread of more severe/contagious forms of an infectious disease^1^. During the SARS-CoV-2 pandemic, national pathogen genome sequencing programs have been substantially enhanced allowing the systematic identification of novel SARS-CoV-2 lineages associated with distinct epidemiological events and geographic regions. At the time of writing [03/06/2021] more than 1.8 million SARS-CoV-2 genomic sequences from 181 distinct countries are incorporated in the GISAID database^2^.

Pango, a sophisticated phylogenetic classification system developed by Rambaut et al^3^, has been endorsed by international health authorities and currently represents the gold standard for the classification of SARS-CoV-2 viral lineages^4^. This system implements a hierarchical four-level nomenclature. Distinct lineages are established based on phylogenetic evidence and important epidemiological and biological events to the evolution of emerging/novel viral strains. Pango delineates a total of 1290 distinct SARS-CoV-2 lineages. Among these, 16 lineages are currently under scrutiny by international health authorities due to their widespread and sustained circulation, coupled with the presence, in the spike glycoprotein, of recurrent mutations associated with increased infectivity and/or reduced neutralization by antibodies^5,6^. Collectively these lineages are known under the acronym VOC (Variant of Concern) or VOI (Variant of Interest) and represent a substantial source of concern for the success of national vaccination campaigns and the effectiveness of measures for the containment of COVID-19^7^. SARS-CoV-2 VOCs and VOIs show a highly biased geographic distribution (Table 1), and until recently^8^ were typically designated by the country of first isolation. While most lineages are compact in size (median 91.5 distinct genomes), the 10 most numerous lineages in the Pango nomenclature incorporate more than 67% of the available genomes. For example, the B.1.1.7 VOC lineage (also known as 20I/501Y.V1 or Variant of Concern 202012/01 (VOC-202012/01)), accounts for over 795 thousand genomes, and is currently the most widely sampled in Europe and worldwide. It was first detected in southern UK in mid-December 2020, and by the beginning of January 2021 had become the most prevalent lineage in that country^9^, with similar dynamics subsequently observed in other European countries^10^. Similar to other VOCs, specific mutations in the B.1.1.7 lineage have been linked with an increased ability to escape from neutralising antibodies and a higher affinity for the human viral receptor hACE2^11^. Data from the UK and Ireland suggest that B.1.1.7 exhibits increased transmissibility^9^, and preliminary observations also suggest a greater disease severity and higher mortality rates^12^.

**Table 1:**
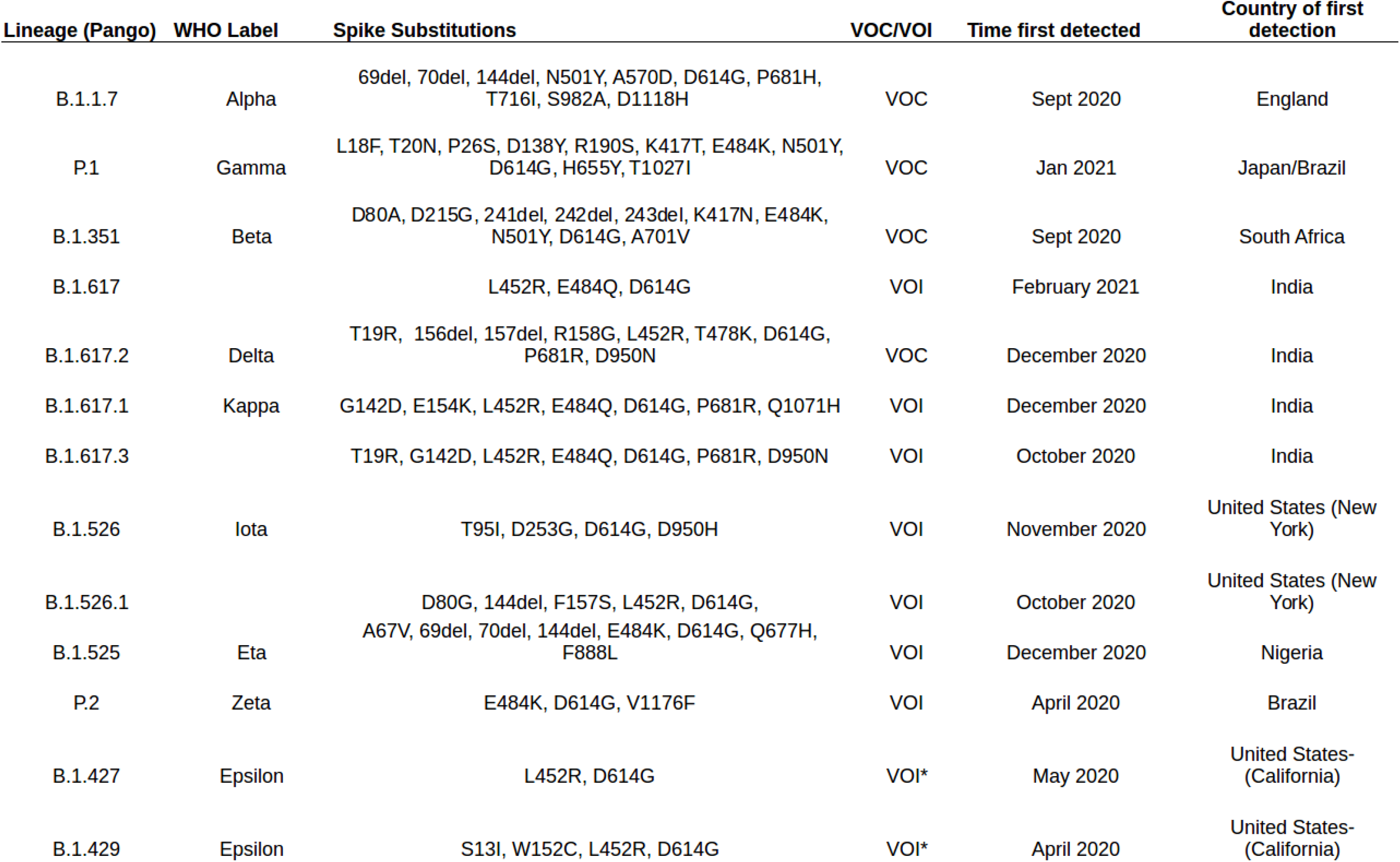
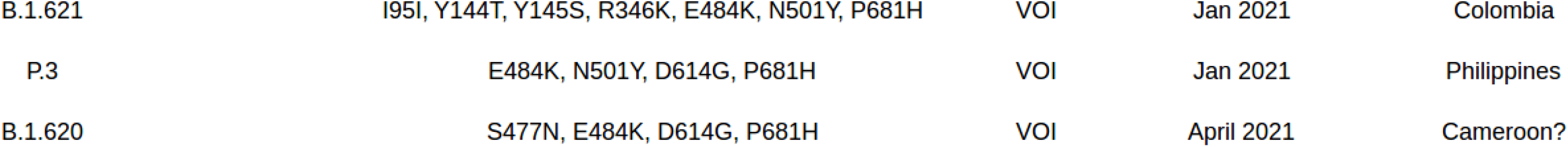
List of Pango lineages currently recognized as VOC/VOI. Lineage (Pango): lineage name according to the pango nomenclature. Substitutions Spike: list of characteristic non-synonymous substitutions in the spike glycoprotein. VOC/VOI: status, VOC or VOI. Time first detected: time of isolation of the first genome associated with the lineage. Country of first detection: country where the lineage was originally isolated.

Although the widespread circulation of B.1.1.7, at least in Western countries, reflects the evolutionary success of this lineage and the efficacy of contemporary approaches to SARS-CoV-2 genomic surveillance in tracking VOCs, several lines of evidence suggest that current sampling of SARS-CoV-2 genomic sequences might be substantially biased^13^ due to various factors including: local incidence of COVID-19^14^, access to state of the art molecular diagnostic test and sequencing facilities^15,16^, bioinformatics bottlenecks in the analyses of the data and sharing of the results^17,18^, as well as the application of different rationales and criteria by local health authorities in the implementation of genomic surveillance strategies. These biases pose relevant questions about the accuracy and completeness of current estimates of the prevalence of SARS-CoV-2 lineages, with possible implications for genomic surveillance of the pathogen.

We and others^19,20^ have recently proposed strategies based on the phenetic clustering of prevalent alleles (with Allele Frequency, AF, > 1%) for the delineation of current and emerging viral genetic diversity in SARS-CoV-2. Here we extend our approach to integrate regional estimates of allele frequency and to facilitate a more rapid and accurate delineation of local variants of SARS-CoV-2. By applying our revised approach to the complete collection of more than 1.8M complete genome sequences of SARS-CoV-2 we demonstrate that our method can identify all current VOCs and VOIs in an unsupervised manner, with reduced turn-around times and high accuracy. Additionally, and particularly in the large and rapidly spreading B.1.1.7 lineage, we outline the presence of several genetic variants with highly biased geographical distributions, which are not captured by current nomenclature standards and which represent a hidden layer of genomic diversity. Finally, by applying simple unsupervised learning methods, based on state of the art clustering algorithms and dimensionality reduction techniques, we present a simple, yet accurate computational strategy for the rapid and automatic identification of VOC and VOI-like SARS-CoV-2 lineages, and provide interesting observations concerning the recurrent mutations associated with these viral variants.

Taken together, we believe that the main results and the approach outlined in this work can provide a substantial contribution to current and future methods for genomic surveillance strategies of human pathogens. A collection of tools and software for the automated analysis of SARS-CoV-2 genomic sequence data is made publicly available at https://github.com/matteo14c/assign_CL_SARS-CoV-2.

## Results

### Biased sampling and geographic diversity in SARS-CoV-2 genomic data

Currently, submissions of publicly available SARS-CoV-2 genomic sequences from the USA and the UK account for more than 65% of currently available genomic sequences. As outlined in Supplementary Figure S1A, the total number of deposited genomic sequences is extremely variable across different geographic regions. Moreover, for many countries, sequence production does not correlate with the incidence of COVID-19 (Supplementary Figure S1B). Indeed, the large majority of SARS-CoV-2 lineages currently identified by the Pango nomenclature are observed only within specific geographic regions (Supplementary Figure S2A), and more than 708 lineages (55% of the total number) reach a overall prevalence of 1% or above only in North America and/or Europe. Accordingly, a limited correlation is observed between the incidence of COVID-19 and the number of Pango lineages identified in different countries (Supplementary Figure S2B)

A total of 104,877 distinct genetic variants were recovered in the complete collection of SARS-CoV-2 genomic sequences considered in this study (Supplementary Table S1). Analyses of allele frequency distributions were performed on aggregated data, on overlapping intervals of 10 days starting from December 26th 2019, i.e., the reported day of isolation of the first sequenced SARS-CoV-2 genome. Consistent with previous reports^19^, only a limited number of variants (1,261, 1.2%, see “HF_cumulative” values in Supplementary Table S2) exhibited a prevalence of 1% or above in the viral population at more than one time interval, while the large majority (103,616, 98.8%) never reached this minimum frequency threshold. Strikingly, when genomic sequences were partitioned according to their respective macro-geographic regions (as defined inSupplementary Table S2), equivalent analyses revealed a more than 7 fold increase in the number of high frequency alleles (from 1,261 to 10,014; compare “HF_cumulative” to “HF_regional” values in Supplementary Table S2), consistent with the observed geographic sampling bias. Dimensionality reduction analyses of allele frequency distribution performed on the complete collection of 104,877 genetic variants at three time intervals (Figure 1A, B, C), demonstrated clear separation between distinct geographic macro-areas irrespective of the time interval considered, with the European and African lineages showing the highest degree of separation on both axes. Equivalent patterns were observed when single countries were considered (Supplementary Figure S3A, B, C). Interestingly, a more clear separation is observed between European and non-European macro-areas (and countries) starting from approximately day 350 (Figure 1C and Suppl. Figure S3C), i.e., in the interval of time that corresponds with the rapid fixation of the VOC B.1.1.7 in many European countries^21^.

**Figure 1:**
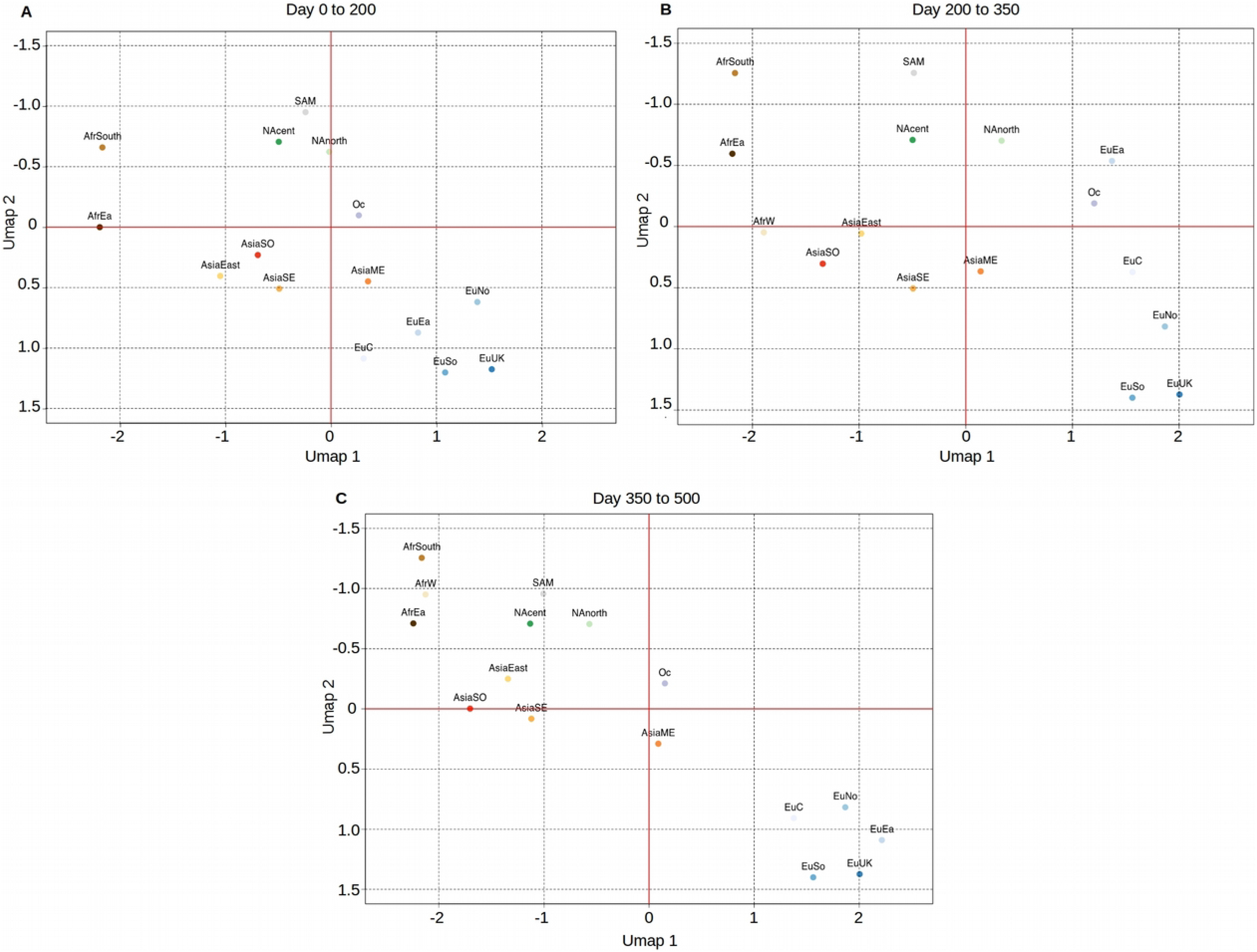
Principal component analysis of local allele frequencies in different geographic areas. Three distinct time intervals are considered, with time T0 set at Time 0= December 26th 2019, i.e., the day of reported isolation of the first SARS-CoV-2 genomic sequence. A: from day 0 to 200. B: from day 200 to 350. C: from day 350 to 500. The following acronym are used for different geographic areas: AfrSouth: Southern Africa; AfrW: Western Africa; AfrEA: Eastern Africa; SAM: South America; NAcent: central America; NAnorth: North America; AsiaEast: Eastern Asia; AsiaSO: Southern Asia; AsiaEast: Eastern Asia; AsiaSE: South Eastern Asia; Oc: Oceania; AsiaME: Middle East; EuC: Central Europe; EuNo: Northern Europe; EuEa: Eastern Europe; EuSo: Southern Europe; EuUK: United Kingdom. See Table S1 for the correspondence between geographic areas and countries.

Notably, a total of 14,484 distinct alleles, corresponding to 13.8% of the total number, were found to display a significant bias in their geographic distribution (Benjamini corrected Fisher test p-value ≤ 0.01, Supplementary table S3). Additionally, when genomic assemblies submitted from different countries were compared we also noticed that genome assembly size distributions (Supplementary Figure S4) differed significantly between submitting countries, resulting in a possible, additional source of technical bias. Indeed, when the original criteria established in Chiara et al^20^ for the definition of “high quality” assemblies were applied, large discrepancies - from 97.6% (Turkey) to 49.2% (Brazil) - were observed in proportions of “high quality” genomic sequences at national level (Supplementary Table S4).

We reasoned that systematic differences in the number of genomic sequences available from distinct geographic areas, together with the observed discrepancies in their overall quality and completeness, could represent relevant issues for the correct classification and mapping of SARS-CoV-2 genomic diversity. To investigate this hypothesis we decided to modify our recently developed^19^ system for the classification of SARS-CoV-2 genomic sequences by:

1. relaxing the criteria for the definition of “high quality” genomic sequences (originally defined as sequences longer than 29,850 nt and including less than 150 ambiguous sites), in order to obtain a more uniform sampling;
2. incorporating regional estimates of allele frequencies;
3. refining the criteria for the definition of high frequency alleles (originally defined as those achieving an allele frequency greater than 1% for more than 50 days in total).

### Modifications of these three classification criteria were as below

1. Given the potential for affecting estimates of allele frequencies, and the observation that the large majority (87%) of “low quality” sequences were incomplete either at 5’ and/or 3’ UTR, criteria for inclusion of sequences in our analyses were partially relaxed (see Materials and Methods). As a consequence, a more uniform proportion of “high quality” genomes was observed across different countries (see Supplementary Table S4).
2. Allele frequencies were estimated separately for different geographic areas (as defined in Suppl Table S1), using time intervals of 10 days, overlapped by 5 days, similar to the approach used in Chiara et al^19^. For every macro-area, only time points associated with at least 100 genomes were considered.
3. Different values for the definition of high frequency alleles were tested, using both different allele frequency thresholds (0.5 to 5%) and different thresholds for their persistence in the viral population (10 to 100 non-consecutive days). The different combinations were evaluated both with respect to the total number of haplogroups (HG) formed, and considering their ability to reconstruct groups corresponding to the current 4 VOCs and 12 VOIs in a timely and accurate manner. The results of these tests are summarized in Table 2. A minimum allele frequency of 1% and a persistence threshold of 15 or more days were selected as the most appropriate combination of parameters for the definition of high-frequency alleles in our classification system. When applied to the complete collection of 1.8M SARS-CoV-2 genomic sequences, the selected combination of parameters resulted in the delineation of a total of 399 distinct HGs, compared with the 238 HG that would be formed by applying our original criteria. Importantly, while defining a relatively compact number of distinct haplogroups, the selected setting was still capable of assigning each of the 16 current VOC/VOI to at least one distinct designation. As reported in Table 3, all the VOC/VOI related groups were formed in an unsupervised manner within a maximum of 20 days from the emergence of the first genome (average 20). This interval of time is comparable to or shorter than that required for their acknowledgment/recognition by international health authorities.

**Table 2:** Combinations of parameters used in the optimization of phenetic clustering of SARS-CoV-2 genomes. AF threshold: threshold of minimum allele frequency. Persistence threshold: number of days that an allele occurs above the minimum AF threshold. VOC HGs: number of VOC lineages associated with at least 1 distinct HG compared to the total VOC. VOI HGs: number of VOI lineages associated with at least 1 distinct HG compared to the total VOI. Tot HGs: Total number of HGs identified. Subsequent columns (Europe, Asia, N. America, S. America, Africa, Oceania) report the total number of HGs formed in distinct continents. The optimal combination of parameters is indicated in red and bold.

| AF threshold | Persistence threshold | VOC with > 1 HGs / total VOC | VOI with > 1 HGs/ total VOI | Tot HGs | Europe | Asia | N. America | S. America | Africa | Oceania |
| --- | --- | --- | --- | --- | --- | --- | --- | --- | --- | --- |
| 0.5 | 10 | 4/4 | 12/12 | 2553 | 1378 | 104 | 1046 | 72 | 30 | 382 |
| 0.5 | 15 | 4/4 | 12/12 | 2346 | 1266 | 95 | 961 | 62 | 24 | 350 |
| 0.5 | 25 | 4/4 | 12/12 | 1689 | 911 | 68 | 691 | 60 | 22 | 253 |
| 0.5 | 50 | 3/4 | 9 | 1123 | 606 | 45 | 459 | 39 | 22 | 168 |
| 0.5 | 75 | 3/4 | 8 | 1017 | 548 | 40 | 416 | 36 | 19 | 151 |
| 0.5 | 100 | 2/4 | 5 | 815 | 440 | 33 | 333 | 34 | 15 | 122 |
| 1 | 10 | 4/4 | 12/12 | 821 | 443 | 33 | 336 | 28 | 15 | 123 |
| **1** | **15** | **4/4** | **12/12** | **399** | **217** | **16** | **165** | **14** | **7** | **60** |
| 1 | 25 | 4/4 | 10/12 | 269 | 145 | 11 | 110 | 8 | 4 | 39 |
| 1 | 50 | 4/4 | 9/12 | 238 | 128 | 8 | 96 | 7 | 4 | 35 |
| 1 | 75 | 3/4 | 8/12 | 221 | 118 | 8 | 90 | 7 | 4 | 33 |
| 1 | 100 | 2/4 | 5/12 | 203 | 108 | 7 | 82 | 6 | 3 | 29 |
| 2.5 | 10 | 4/4 | 9/12 | 375 | 202 | 14 | 152 | 11 | 15 | 56 |
| 2.5 | 15 | 4/4 | 7/12 | 326 | 176 | 13 | 133 | 9 | 11 | 48 |
| 2.5 | 25 | 4/4 | 7/12 | 259 | 139 | 9 | 105 | 8 | 6 | 38 |
| 2.5 | 50 | 2/4 | 6/12 | 221 | 118 | 8 | 90 | 7 | 4 | 33 |
| 2.5 | 75 | 2/4 | 5/12 | 203 | 108 | 7 | 82 | 6 | 3 | 29 |
| 2.5 | 100 | 2/4 | 3/12 | 161 | 86 | 6 | 66 | 34 | 2 | 24 |
| 5 | 10 | 4/4 | 7/12 | 212 | 114 | 7 | 86 | 6 | 3 | 30 |
| 5 | 15 | 2/4 | 4/12 | 182 | 97 | 6 | 74 | 5 | 3 | 26 |
| 5 | 25 | 2/4 | 3/12 | 133 | 71 | 4 | 53 | 4 | 2 | 19 |
| 5 | 50 | 1/4 | 2/12 | 117 | 62 | 4 | 47 | 3 | 2 | 17 |
| 5 | 75 | 1/4 | 1/12 | 111 | 59 | 4 | 45 | 3 | 2 | 16 |
| 5 | 100 | 1/4 | 1/12 | 107 | 57 | 4 | 44 | 3 | 1 | 15 |

**Table 3:** Time from emergence of a VOI/VOC to formation of an HG in our system. Lineage (Pango): lineage according to the Pango nomenclature. VOC/VOI: status, VOC or VOI. Time first detected: time of isolation of the first genome associated with the lineage. Days from first detection to HG: days intercurring from the isolation of the first genome to the appearance of a corresponding HG. Days from detection to VOC-VOI: days intercurring from the isolation of the first genome to identification of a VOC/VOI

| **Lineage (Pango)** | **VOC/VOI** | **Time first detected** | **Days from first detection to HG** | **Days from detection to VOC-VOI** |
| --- | --- | --- | --- | --- |
| B.1.1.7 | VOC | October 2020 | 15 | 63 |
| P.1 | VOC | December 2020 | 18 | 21 |
| B.1.351 | VOC | Sept 2020 | 17 | 54 |
| B.1.617 | VOI | February 2021 | 21 | 31 |
| B.1.617.1 | VOI | December 2020 | 22 | 59 |
| B.1.617.2 | VOC | December 2020 | 21 | 59 |
| B.1.617.3 | VOI | October 2020 | 21 | 59 |
| B.1.526 | VOI | November 2020 | 18 | 45 |
| B.1.526.1 | VOI | "October 2020" | 19 | 45 |
| B.1.525 | VOI | December 2020 | 21 | 35 |
| P.3 | VOI | January 2021 | 20 | 35 |
| B.1.427 | VOI | May 2020 | 23 | 291 |
| B.1.429 | VOI | July 2020 | 19 | 64 |
| B.1.621 | VOI | Jan 2021 | 18 | 23 |
| P.2 | VOI | Jan 2021 | 23 | 21 |
| B.1.620 | VOI | April 2021 | 24 | 35 |

### Hidden genomic diversity in large SARS-CoV-2 lineages

Applied to the 1.8 Million SARS-CoV-2 genomic sequences available in the GISAID database on 3th June2021 (Supplementary Table S5), our method delineated a total of 399 distinct HGs in an unsupervised manner. We identified several distinct HGs whose merging in the canonical nomenclature appears due to saturation of its four level system.

Strikingly, 144 HGs fall within the B.1.1.7 lineage: HG26, which is defined by the same set of alleles as the original designation of the lineage, and 143 additional haplogroups all of which show a distinctive pattern of high frequency alleles and a distinct geographic distribution (Supplementary Table S6). This observation suggests a widespread unexplored diversity in the genomes associated with this lineage. Based on the analysis of the first 50 isolates for each of these 144 HGs (isolation dates as reported in GISAID) (Supplementary Table S7) and on their geographic distribution (Supplementary Table S8), 46 HGs, including HG26, were associated with the UK, while 24 HGs were preferentially observed in the USA. A limited, but consistent number of HGs were associated with distinct European countries, including: Germany (6 HGs), Sweden (9 HGs), Denmark (3HGs), Italy (2 HGs), the Netherlands (2 HGs), Norway, Finland Belgium and Estonia (1HG each). The remaining 42 B.1.1.7 associated HGs could not be tentatively associated with a specific country due to their more variable geographic distributions.

As reported in Supplementary Table S6, a total of 345 high frequency variants (AF > 1%) are associated with the 144 HGs corresponding to the B.1.1.7 lineage. Each of these haplogroups is itself defined by c.a. 32 (average 32.5) distinct high frequency alleles. Interestingly, a highly significant positive correlation is observed between the time of first isolation and the total number of high frequency variants associated with each HG (Pearson correlation 0.8423, p-value < 2.2e-16), suggesting that the emergence of novel HGs probably results from the a continuous fixation of novel variants in a pre-existing genomic background. Cumulatively, 26 markers are common to more than 142 HGs, while the large majority (302) are associated with 5 or less HGs.

In general HGs associated with B.1.1.7 display a higher number of “defining” markers compared to the remaining HGs (32.5 compared to 18.8 on average), however collectively they do not display increased levels of intra-group variability (Figure 2). Consistent with this observation, when “defining” high frequency variants are excluded, an overall similar proportion of sites potentially associated with positive selection, as determined by Hyphy^22^, is recovered between the 2 groups (Supplementary Table S9). Taken together these observations suggest that the “increased” diversity associated with Pango lineage B.1.1.7 likely reflects the widespread circulation of this virus strain coupled with a more capillary sampling of SARS-CoV-2 genomes from European and Western countries in general, rather than an alteration of the evolutionary dynamics of the strain.

**Figure 2:**
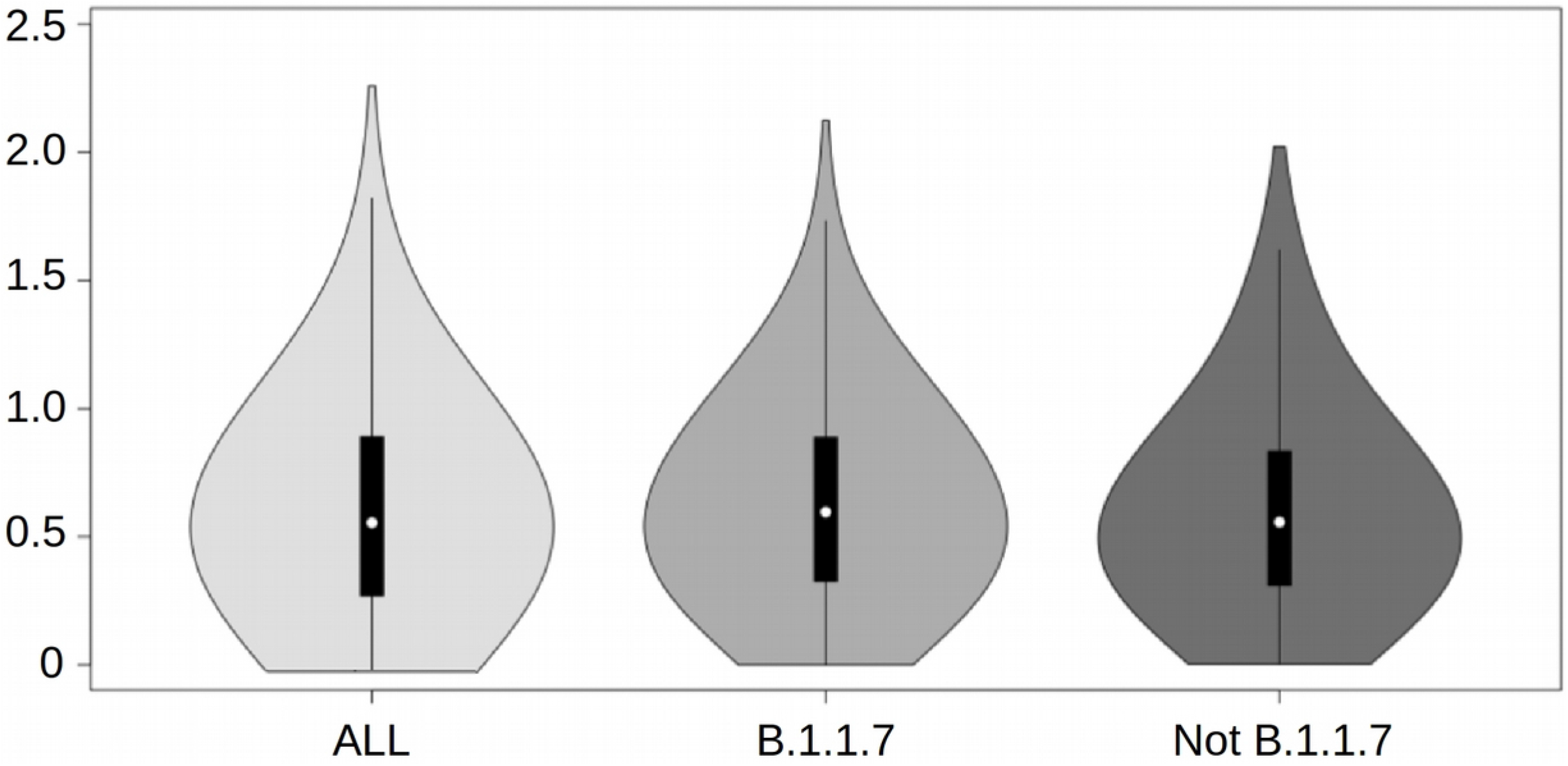
Distribution of genetic distance, computed as the total number of distinct polymorphic sites between closest pairs of genomes. Distributions are represented in the form of a violin plot. Distances are indicated on the Y axis.

### Functional evaluation of high frequency variants fixed in distinct HGs

Although cumulatively the majority of the high frequency variants identified by our method within the B.1.1.7 lineage are not currently associated with mutations of recognized epidemiological relevance, several notable exceptions were observed.

HG85, HG166, HG214 and HG215, which collectively incorporate over 10,500 genomic sequences, are partly defined by a c.52C>T mutation (L18F in the S spike glycoprotein). L18F is currently observed in 8.8% of all available SARS-CoV-2 genome sequences (Supplementary Table S6) and has rapidly become established from independent origins, including in HG21, a large haplogroup related to/descending from the B.1.177 lineage which rapidly spread across Europe in the summer of 2020^22^, and in multiple HGs which correspond to Pango P.1 lineage (HG298, and HG365 and HG389-HG392), and the B.1.351 lineage (HG161, HG349 and HG397). Both P.1 and B.1.351 are classified as VOCs by international health authorities due to their increased infectivity and capacity to escape neutralising antibodies^7^. Notably, L18F is adjacent to a glycosylation site and has previously been associated with escape from several antibodies that target the N-terminal domain (NTD) of the S protein^24^.

Similarly, HG284 - a small HG consisting of c.a. 900 genomic sequences which is specifically associated with Sweden - 79.4% of the genomes forming this HG being deposited from that country (Supplementary Table S8) - includes the S494P substitution in the recognition binding motif of the spike glycoprotein. This substitution is predicted to be under positive selection according to Hyphy, has been recently shown to increase the affinity for the host target receptor hACE2^25^, and has been postulated to facilitate the escape from the recognition by some antibodies^26^. Currently, B.1.1.7+S494P is recognized as a “distinct” (from B.1.1.7) VOI by international health authorities, and has been subjected to more dedicated monitoring.

Unsurprisingly, the ORF8 gene, which is thought to have been inactivated by a stop-gain in an ancestor of the B.1.1.7 lineage^27^, is associated with an excess of deleterious substitutions in HGs corresponding with the B.1.1.7. These include: a further stop-gain mutation at residue K68, which is collectively associated with 45 HGs and more than 340K genomic sequences; a single nucleotide deletion (at residue I121) which is associated with more than 27K sequence and 6 HGs; and a single base insertion (at residue 119D) which is observed in 4 HGs and affects more than 22K genomes (Supplementary Table S6). Both of these indels would cause the elongation of an intact ORF8 by a few amino acids (I121SK and DFI119EFHLN respectively in Supplementary Table S6).

Additionally, we observe that 14 HGs, with a total of more than 64K genomic sequences and c.a. 9% of the viral isolates currently associated with B.1.1.7, carry a premature truncation of the Orf7a protein (184C>T,p.Q62*). Interestingly, several independent studies have reported SARS-CoV-2 isolates with unique in- and out-of-frame deletions in Orf7a resulting in heavily truncated and/or non functional polypeptides^28,29^. Although the implications of these mutations are not completely understood at present, we observe that more recently, depletion of Orf7a has been linked with reduced inactivation of the host immune system by SARS-CoV-2^30^.

A total of 2,762 genomes constituting HG114 display all of the single nucleotide polymorphisms associated with B.1.1.7, but lack the 3 deletions that also define this lineage^27^ (spike:IHV68I and YY144Y; nsp6:SGF106 in Supplementary Table S6). Although it might be tempting to speculate that this haplogroup could represent an early divergence from B.1.1.7, pre-dating the deletions, the first genome of this type was sampled more than 100 days after the first B.1.1.7 genome. Accordingly, we tested the node purity (see Materials and Methods) of HG114 on the SARS-CoV-2 genome tree as obtained from the Audacity tool in GISAID, and noted a significantly (56.56%) lower degree of phylogenetic coherency (Wilcoxon p-value ≤2e-16) for this group compared to other haplogroups defined by our method (mean 82.58, Supplementary Table S10). HG114 is also associated with a reduced percentage of “high quality” (according to our original definitions) genome assemblies (63.76%, compared with an average of 82.28%). Taken together with the patchy geographical distribution of HG114 (Supplementary Table S7 and S8), we speculate that the apparent HG114 grouping results from clustering of “artifactual” sequences derived from systematic errors in variant calling pipelines that lack adequate indel calling capacity.

Outside of the B.1.1.7 lineage, we observe that 25 additional Pango-defined lineages with large numbers of genomes (Table 4) correspond to multiple HGs in our system. Cumulatively, these lineages are linked to 105 different HGs. Strikingly, this list includes all the current VOCs: B.1.1.7, P.1, B1.617.2 B.1.351 and two VOIs: B.1.429 and B.1.526 (P-value Fisher for VOCs/VOIs over-representation=0.001351).

**Table 4:** List of lineages corresponding with multiple HGs. Lineage (Pango): lineage according to the pango nomenclature. Tot HGs: Total number of HGs associated with the lineage. Is VOC/VOI?: is the lineage a VOC/VOI: 1=TRUE, 0=FALSE. Tot genomes: total number of genomes in the lineage. List HGs: list of HGs associated with the lineage

| Lineage<br>(Pango) | Tot HGs | Is VOC/VOI ? | Tot genomes | List HGs |
| --- | --- | --- | --- | --- |
| B.1.1.7 | 144 | 1 | 800127 | 26 76 77 78 79 80 81 84 85 103 104 114 137 138 139 140 141 142 143 144<br>145 146 147 148 154 155 163 164 165 166 168 169 171 172 173 211 212 213<br>214 215 216 217 218 219 220 221 222 223 224 225 226 227 228 229 230 231<br>232 264 265 266 267 268 269 270 271 272 273 274 275 276 277 278 279 280<br>281 282 283 284 285 286 287 288 289 290 291 292 293 294 295 296 297 299<br>300 301 302 303 304 305 306 307 308 327 328 329 330 331 332 333 334 335<br>336 337 338 339 340 341 342 345 346 347 350 351 352 353 354 355 356 357<br>358 362 363 364 373 374 375 376 377 378 379 380 381 382 383 384 |
| B.1.177 | 10 | 0 | 57121 | 21 53 55 100 116 134 201 203 204 252 |
| B.1.1.214 | 8 | 0 | 17686 | 108 111 112 119 313 314 315 318 |
| B.1.2 | 7 | 0 | 93377 | 23 58 95 96 136 151 326 |
| D.2 | 7 | 0 | 12848 | 15 190 191 192 193 194 195 |
| B.1.1.37 | 6 | 0 | 5018 | 20 28 50 51 200 236 |
| B.1.160 | 6 | 0 | 24354 | 24 48 49 106 247 312 |
| B.1.177.21 | 6 | 0 | 13010 | 61 258 259 260 261 262 |
| B.1.221 | 4 | 0 | 13376 | 25 59 64 65 |
| B.1.429 | 4 | 1 | 31369 | 87 157 158 348 |
| B.1.1.284 | 3 | 0 | 9126 | 99 309 310 |
| B.1.177.12 | 3 | 0 | 6651 | 60 256 257 |
| B.1.177.45 | 3 | 0 | 2904 | 57 253 254 |
| B.1.279 | 3 | 0 | 2121 | 82 83 105 |
| B.1.526 | 4 | 1 | 22029 | 152 170 343 386 388 |
| AD.2 | 2 | 0 | 4427 | 22 207 |
| B.1.1.519 | 2 | 0 | 19282 | 159 160 |
| B.1.177.65 | 2 | 0 | 1447 | 205 206 |
| B.1.234 | 2 | 0 | 6373 | 118 317 |
| B.1.243 | 2 | 0 | 11181 | 68 72 |
| B.1.258 | 2 | 0 | 8801 | 27 234 |
| B.1.351 | 2 | 1 | 23958 | 161 349 387 |
| B.1.497 | 2 | 0 | 2990 | 31 88 |
| B.1.596 | 2 | 0 | 4718 | 208 209 |
| P.1 | 6 | 1 | 29183 | 298 365 389 390 391 392 |
| B.1.617.2 | 7 | 1 | 15315 | 367 393 394 395 396 397 398 |

For example, while the “original” designation of the P.1 VOC corresponds to HG365 in our system, we observe (Supplementary Table S11) the presence in P.1 of five additional HGs, i.e., HG298, HG389-HG392, which collectively carry a total of 16 additional high frequency variants, including four distinct non synonymous substitutions predicted to be under positive selection (Supplementary Table S9) according to Hyphy^23^. Similarly, the B.1.351 lineage is associated with three distinct HGs (Supplementary Table S12) in our system: HG161, which is defined by the same set of alleles included in the original Pango designation of this lineage, HG349, which carries 4 additional high frequency mutations, including 2 non-synonymous substitutions in the nsp2 (V649F) and nsp12 (Q822H) genes, respectively, and HG387 which harbors an allele of the orf3a gene inactivated by a stop-gain mutation.

The B.1.617.2 VOC, which is rapidly becoming the most prevalent variant of SARS-CoV-2 in the UK, is associated with 8 distinct HGs (HG367, and HG393-HG399) in our classification system (Supplementary Table S13). While all the HGs share the unique combination of 28 high frequency variants that define this lineage, a total of 76 distinct variants are observed within B.1.617.2 according to our analyses of which 21 are specific only to a single HG. Of the 10 missense substitutions associated with a single HG in B.1.617.2, 8 are predicted to be under positive selection (p-val Fisher 0.0003891). Interestingly this list includes the A222V missense substitution in the spike glycoprotein (HG394), a mutation that is considered to be characteristic of the B.1.177 lineage which rapidly spread across Europe in the summer of 2020^22^.

Interestingly, additional HGs associated with VOC/VOI lineages are often observed only in the countries/geographic regions where the corresponding VOC/VOIs were originally isolated. For example HGs corresponding with the B.1.526 and B.1.429 VOIs are highly prevalent in North America, while HGs corresponding with B.1.617.2 are highly prevalent in India and in the UK. However we also notice a limited number of HGs with a more widespread geographic distribution, which are likely to reflect independent introduction of the same HGs in distinct countries; these include some HGs associated with the P.1 and B.1.351 VOCs, like, for example, HG161 and HG387 (B.1.351) and HG390 and HG365 (P.1).

Globally, the 26 aforementioned lineages associated with multiple HGs include about 67% (Table 4) of all the 1.83M genomic sequences considered in our analyses, further emphasizing the extent of “emerging high frequency variants” not explicitly addressed by current nomenclature standards.

Taken together, these observations highlight the importance of the incorporation of local estimates of allele frequency into systems for the genomic surveillance of SARS-CoV-2 for more refined tracking of the routes of circulation of variants across and within different countries.

### A simple approach for the prioritization of SARS-CoV-2 variants

Simple descriptive features (Supplementary Table S14), including total number of synonymous and non-synonymous variants, measures of global genetic diversity, and counts of sites predicted to be under positive and/or negative selection, were recorded for all of the 1290 Pango lineages and the 399 distinct HGs defined in this study. Counts were normalised using unity based normalization, and the Louvain clustering algorithm^31^ was applied to identify groups of HGs or Pango lineages with similar profiles of mutations. Based on different cluster stability metrics, 8 and 3 clusters were identified as the optimal clustering solution for the Pango lineages and the HGs, respectively. The dimensionality reduction plots obtained with Umap^32^, and reported in Figure 3A and 3B, show a good level of separation between the clusters. Strikingly, in both Figure 3A and 3B we observe that the large majority of VOCs/VOIs grouped in a single cluster: in the analysis of the Pango lineages, 15 over 16 VOCs/VOIs grouped in cluster 2 (see C2lin in Figure 3A); in the analysis of HGs, with the only exception of HG114 (i.e., the HG in B.1.1.7 that might be associated with systematic sequencing/assembly errors and was assigned to cluster 1 (see C1HG in Figure 3B), all the HGs corresponding to the 16 VOCs/VOIs were contained in cluster 3 (see C3HG in Figure 3B). These results clearly indicate that the simple metrics adopted in this work can reliably identify compact and discrete groups of SARS-CoV-2 variants that incorporate all current VOCs/VOIs regardless of the classification system (Pango or HGs). This suggests that they could both be effectively applied to facilitate the identification of potential VOCs and VOIs emerging in the future. Although the equivalence between the HGs of our system and Pango is not complete, cumulatively high levels of agreement were observed between C2lin and C3HG, with more than 67% of the Pango lineages/HGs being common between the two clusters (P-val hypergeometric ⩽ 1 e-16). Compared to the remaining clusters, clusters associated with VOCs and VOIs displayed an overall increase in genomic diversity, a higher number of substitutions (both synonymous and nonsynonymous), and an increased proportion of incompletely fixed alleles, irrespectively of the allele frequency threshold (see Supplementary Figure S5A and B). This notwithstanding, levels of intra-group genomic diversity were similar across HGs and lineages assigned to different clusters, suggesting that the overall increase in diversity in the HGs and lineages associated with VOCs and VOIs does not reflect an accelerated evolutionary rate (Supplementary Figure S6).

**Figure 3:**
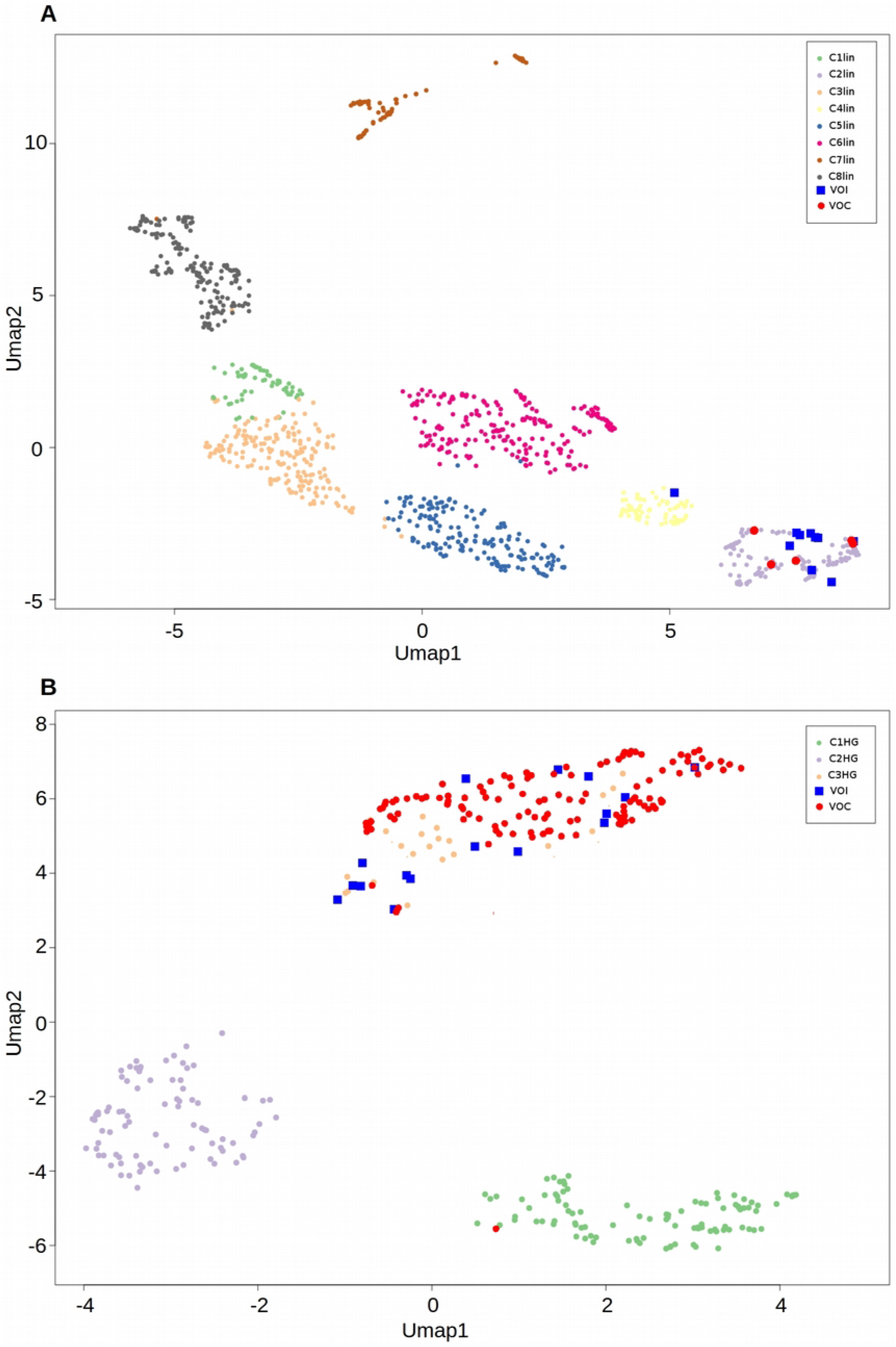
Umap representation of clusters of Pango lineages and HGs. A: Umap of Pango lineages. Distinct colors (see legend) are used to represent distinct clusters. VOC lineages are represented as large solid red dots and VOI lineages as solid blue squares. B: Umap of HGs defined by our system. HGs associated with VOCs and VOIs are represented as solid red dots and solid blue squares, respectively.

A Fisher-exact test was applied to genetic variants showing a biased distribution across the different clusters. A total of 238 and 114 allele variants were found to be over-represented in C2lin and C3HG, respectively (Supplementary Table S15). Consistent with the observed overall increase in genomic diversity in these clusters, equivalent analyses performed in any of the other clusters recovered only a total of 13 over-represented alleles. A total of 93 distinct over-represented alleles were shared by both C3HG and C2lin (see column “common” in Supplementary Table S15; P-val hypergeometric ⩽ 1e-32). Three genomic regions in particular were found to be enriched with mutations over-represented in VOC/VOI containing clusters: the S glycoprotein (30 substitutions in total, of which 26 were non-synonymous and 3 inframe deletions); the N protein (16 substitutions of which 13 missense); and s2m (5 substitutions), a 43 bp secondary structure element of unknown function which is found in several families of single-stranded RNA viruses^33^. Importantly, we observe that of the 84 genetic variants in protein coding genes, 69 are associated with non synonymous substitutions and 21 are predicted to be under positive selection (P-value hypergeometric ⩽ 1e-18); strikingly these include all of the most relevant mutations associated with VOCs and VOIs in the S glycoprotein, including E484K, N50Y and L452R, along with several additional mutations, whose significance has been already confirmed by *in vitro* studies (see for example L18F, G142D, S494P and others mutations in Supplementary Table S15).

## Discussion

Genomic surveillance of SARS-CoV-2 requires rapid and effective approaches for the identification of novel strains/lineages as they emerge, coupled with accurate models for the prioritization of variants of potential concern and the identification of strains that should be subjected to more careful/strict observations and characterization. Ideally these approaches should be completely automated, to reduce turn-around times and to avoid potential biases in the classification of viral strains/lineages.

Here, we use a simple and transparent analysis of the available SARS-CoV-2 genomic sequences, to demonstrate systematic and recurrent biases in the current sampling of SARS-CoV-2 genomic diversity, and the consequent implications for the rapid characterization of current and future VOCs and VOIs. Additionally, we highlight possible limitations of Pango, the current standard for the classification and nomenclature of SARS-CoV-2, which may prevent the rapid and unsupervised identification of emergent “regional” genomic diversity.

By applying a revised implementation of the strategy proposed in Chiara et al^19^, based on relaxed filters for the inclusion of low-quality genomic assemblies and on the incorporation of regional estimates of allele frequencies, we present a novel, completely automated system for the monitoring of this genomic diversity. Importantly, we observe that our revised approach can correctly associate one or more related haplogroups to all current Variants of Concern (B.1.1.7, P.1, B.1.617.2, B.1.351) and Variants of Interest defined by international health authorities (Table 3). Our approach requires, on average, 20 days between the availability of the first genomic sequence associated with a VOC/VOI and the formation of a corresponding HG. This timeframe is completely in line with that observed for the reporting of current VOCs and VOIs, although no manual intervention is required by our framework.

Since B.1.1.7 is currently the most prevalent and rapidly expanding lineage of SARS-CoV-2, the observation that a substantial proportion of high frequency mutations of in this lineage, emerging at different geographic locations, is not captured by current nomenclature standards, bears important implications for the accurate and rapid tracking of other mutations that will accumulate in this and other genomic backgrounds in the near future. For example, we observe recurrent non-synonymous substitutions in the spike glycoprotein, which are associated with a large number of genomes and have been linked with important functional implications by different studies. Notably, some of these variants are specifically associated with distinct geographic regions (see HG284 in Sweden), suggesting the importance of the incorporation of local allele frequency estimates for the timely identification of relevant new haplotypes. Importantly, we observe that similar considerations apply also to many other rapidly-expanding SARS-CoV-2 lineages (see Table 4), and most important to several VOCs/VOIs (6 over 16). The case of the B.1.617.2 Pango lineage, which is rapidly becoming the most prevalent lineage of SARS-CoV-2 in the UK and which is currently associated with 8 distinct HGs in our system, is emblematic in this respect.

Finally, we show that simple unsupervised clustering approaches of SARS-CoV-2 lineages/haplogroups with similar patterns of mutations can be applied for the identification of SARS-CoV-2 variants of potential epidemiological relevance. Our approach recapitulates several important findings regarding the main mutations in the spike glycoprotein that characterize VOI and VOC lineages, and provides a reliable and automatic method for the prioritization of SARS-CoV-2 variants which might potentially represent a source of concern.

Due to its profound mechanistic importance, and thanks to our detailed appreciation of its molecular interactions, SARS-CoV-2 nomenclature standards have tended to focus on variants in the S gene, whose potential biological significance can often be rapidly flagged through expert intervention. However, the possibility remains that non-spike variants might also affect the epidemiological and clinical course of the pandemic. For example, orf8 disruptions have occurred in several SARS-CoV-2 lineages as well as in later isolates from the 2005 SARS epidemic^34^. While available data hint that orf8 disruption likely reduces disease severity^35^, it is possible, in the context of limited testing and differentially observed self isolation, that “less pathogenic” strains might actually be transmitted more rapidly in some environments. Our finding that approximately 9% of all current SARS-CoV-2 isolates in the B.1.1.7 lineage carry a truncated form of the orf7A protein is also particularly relevant. Indeed, orf7a was recently shown to inhibit IFN-I signaling^30^ while genetic defects in IFN-I signaling in the host have been recently linked with more severe forms of COVID-19^36^. Taken all together these observations might suggest again the potential emergence of less pathogenic/attenuated strains in B.1.1.7.

The approach for the prioritization of potential VOCs/VOIs proposed in this study, as well as for grouping lineages/haplogroups carrying all the currently known epidemiological relevant spike variants in compact and highly consistent groups, recovers an additional set of mutations specifically enriched in SARS-CoV-2 variants with a profile of mutations similar to that of VOCs and VOIs. For example we notice a number of recurrent non-synonymous substitutions in the N nucleocapsid protein. Although the primary functions of the N gene are the binding and packing of the viral RNA genome, N is a highly immunogenic protein and is expressed at very high levels during infection. Interestingly we notice that 6 out of 16 mutations in N over-represented in C2lin/C3HG are associated with positive selection according to Hyphy (Supplementary Table S15) and 9 are linked with predicted epitopes according to Kiyotani et al^37^. While, to the best of our knowledge, none of these mutations is currently considered to be relevant from an epidemiological point of view, these observations might suggest a possible link with immune evasion of anti-N neutralising antibodies, thus warranting further investigations.

Additionally, we observe that 5 alleles of the s2m locus are significantly over-represented in clusters of SARS-CoV-2 variants showing a pattern of mutation similar to VOCs/VOIs, potentially suggesting a possible change in selective pressure. According to recent studies, s2m represents one of the most variable genomic elements in the genome of SARS-CoV-2^19^. Since the reference genome assembly carries a nucleotide substitution at a highly conserved and structurally important position (T 29,758) in s2m, with possible impacts on its structural stability, it was speculated that the observed excess in diversity could be compatible with either loss of functional constraints and/or diversifying selection^20^. Strikingly, we observe that the G to T substitution at 29,742, which is a hallmark of the B.1.617.2 VOC (and related lineages), and is also significantly over-represented in C2lin and C3HG, is associated with a considerable increase in s2m secondary structure stability. Interestingly, with a cumulative frequency of 1.41%, this variant is the most frequent variant in s2m, and shows a highly biased geographic distribution, with a prevalence of 10% and 7% respectively in central and southern Asia. However, the G to C substitution at 29,734 which shows a similar level of prevalence (frequency 1.39%) and a roughly equivalent geographic distribution does not seem to be associated with an increase in stability and is not associated with VOCs/VOIs. Similar considerations apply also to 3 other recurrent substitutions of s2m which are significantly enriched in C3HG and C2lin, but are not associated with VOC/VOI lineages and are predicted to determine only marginal increases in the stability of s2m secondary structure. In the light of these considerations it remains unclear whether these changes reflect ongoing widespread diversifying selection in SARS-CoV-2, or whether the original disruptive substitution inactivated s2m function, leading to a general loss of significant purifying constraints.

In conclusion we believe that the current work provides several distinct lines of evidence to demonstrate the accuracy and efficacy of fully automated “epidemiogenomic” approaches, such as the one described here, in drawing attention to emerging variants of SARS-CoV-2 as epidemiological significant in a timely manner and accurate manner. Based on these considerations we suggest that a more accurate and comprehensive system for the mapping and tracking of SARS-CoV-2 genomic diversity should flank the Pango system.

## Methods

### Data and code availability

A software package for the classification of genomic sequences of SARS-CoV-2 according to the haplogroups defined in the present study is available at: https://github.com/matteo14c/assign_CL_SARS-CoV-2. A standalone Galaxy implementation is available at: http://corgat.cloud.ba.infn.it/galaxy under Tools/utilities for Haplogroup assignment. Extended data with the complete list of genomes analysed in this study and their assignment to HGs is available from the following Figshare repository: https://doi.org/10.6084/m9.figshare.14823366

### Data retrieval

The complete collection of SARS-CoV-2 genomic sequences and associated metadata (including lineage designation according to the Pango nomenclature) was retrieved directly from gisaid.org on June 3rd 2021. Data concerning the incidence and total number of cases of COVID-19 were downloaded from the WHO dashboard, at https://covid19.who.int/ on the same day.

### High quality genomes

Custom Perl scripts were used to infer the size of each genomic assembly and the number of uncalled bases/gaps (denoted by N in the genomic sequence). High quality genomic assemblies were initially defined, according to the criteria outlined in Chiara et al^19^, as genome sequences longer than 29,850 nt and including less than 150 ambiguous sites. Since systematic differences were here observed when comparing genomic assemblies derived from different countries (Table S10), and due to the frequent incompleteness of the 5’- and/or 3’-UTR (having a length of 229 and 265 nt, respectively), these criteria were revised in order to prevent possible biases in the computation of allele frequency distributions. Therefore, sequences with less than 150 ambiguous sites, and with less than 350 bases missing at either the 3’ or 5’ end of the genome (i.e longer than 29,550 nt) were now considered of high quality. Only sequences satisfying these criteria were considered for the computation of allele frequency and for the definition of SARS-CoV-2 haplogroups.

### Haplogroups and functional annotation

Bioinformatics analyses for the delineation of novel HGs and the assignment of genomic sequences to HGs were performed according to the methods described in Chiara et al^20^. Briefly, SARS-CoV-2 genomes were aligned to the 29,903 nt-long reference assembly of SARS-CoV-2 by means of the nucmer program^38^ . Genetic variants were identified by means of the show-snp utility. Allelic frequencies were computed using a custom Perl script. Genomic sequences were assigned to a geographic region and/or a macro-area based on the country of origin of the genome, as reported in the GISAID database. Correspondence between countries and geographic areas is reported in Supplementary Table S1. Functional annotation of genomic variants was performed by CorGAT^39^. Designation of VOC and VOI lineages was in accordance to the ECDC report of June 3rd 2021^40^. Haplogroups were established by hierarchical clustering of phenetic profiles of presence/absence of high frequency alleles, by applying the hclust function from the R standard libraries (Maechler et al, 2019). The cutree function was used to separate distinct clusters at the desired level of divergence (two distinct variant sites).

An haplogroup was considered to be associated with a specific country when 50% or more of the genomic sequences included in the haplogroup were associated with that country, and/or when the same condition applied to the first 50 isolated genomic sequences of the haplogroup.

### SARS-CoV-2 phylogeny, node purity and detection of sites under selection

SARS-CoV-2 phylogeny in newick format was obtained from the Audacity^41^ tool in GISAID^2^ . The “node purity” of SARS-CoV-2 HGs in the tree was established using a custom script based on the ape R package^42^. Briefly, for every HG all the distinct nodes in the tree containing at least 50 genomic sequences of that HG were retrieved, and the proportion of HG members with respect to the total number of sequences descending from the node was calculated. The node purity was computed as the weighted (by the number of genomes) average of the proportions observed at every node. Alignments of SARS-CoV-2 protein-coding genes were performed by the means of the Muscle^43^ software. Identification of sites under selection was performed by applying the MEME and FEL methods, as implemented in the Hyphy package^23^, to the phylogeny and the concatenated alignment of protein-coding sequences. A P-value of 0.05 was considered for the significance threshold. Only sites predicted to be under positive selection according to both methods were considered. For sites predicted to be under negative selection, only FEL was used, since MEME can not identify purifying selection^44^.

### Statistical analyses, dimensionality reduction and clustering

All statistical analyses and tests were performed by means of the stats^45^ R package as provided by the R programming language. Dimensionality reduction analyses were performed by means of the Uniform Manifold Approximation and Projection (Umap) algorithm as implemented in the R umap package^46^. Clustering of mutation patterns of SARS-CoV-2 lineages/HGs was performed by means of the Phenograph algorithm as implemented by the RPhenograph package^47^. The following values were tested for the k (number of nearest neighbours) parameter: 5, 10, 15, 20, 30 and 50. Cluster stability metrics for the evaluation of different clustering solutions were computed by means of the clValid R package^48^.

## Acknowledgments

We thank ELIXIR Italy for providing the computing and bioinformatics facilities and Barbara De Marzo for technical assistance. We gratefully acknowledge the authors, originating and submitting laboratories of the sequences from GISAID’s EpiFlu Database on which this research is based. This work was supported by the Italian Ministero dell’Università e Ricerca: PRIN 2017, Consiglio Nazionale delle Ricerche, H2020 projects EOSC-Life (GA 824087), EOSC-Pillar (GA 857650) and ELIXIR Converge (GA 871075), and Elixir-IIB.

## Supplementary information

**Figure S1:** Total number of genomic sequences and correlation with the incidence of COVID-19. A) total number of genomic sequences deposited at intervals of 10 days from different geographic areas. Time T0 sets at Time 0= December 26th 2019, i.e. the day of reported isolation of the first SARS-CoV-2 genomic sequence. The following acronym are used for different geographic areas: AfrSouth: Southern Africa; AfrW: Western Africa; AfrEA: Eastern Africa; SAM: South America; NAcent: central America; NAnorth: North America; AsiaEast: Eastern Asia; AsiaSO: Southern Asia; AsiaEast: Eastern Asia; AsiaSE: South Eastern Asia; Oc: Oceania; AsiaME: Middle East; EuC: Central Europe; EuNo: Northern Europe; EuEa: Eastern Europe; EuSo: Southern Europe; EuUK: United Kingdom. B) correlation between incidence of Covid-19 (log10, Y axis) and total number of SARS-CoV-2 genomic sequences deposited in public databases (log 10, x axis) in 100 countries for which at least 500 genomes of SARS-CoV-2 have been deposited.

**Figure S2**: A Barplot of total number of Pango lineages by geographic area. The barplot reports the total number of HGs that reach an allele frequency of at least 1% in different geographic areas. Acronyms and color code according to Fig S1. B) correlation between incidence of Covid-19 (log10, Y axis) and total number of Pango lineages deposited in public databases (log 10, x axis)

**Figure S3**: Principal component analysis of local allele frequencies in different countries. Three distinct time intervals are considered, with time T0 set at Time 0= December 26th 2019 i.e., the day of reported isolation of the first SARS-CoV-2 genomic sequence. A: from day 0 to 200. B: from day 200 to 350. C: from day 350 to 500.

**Figure S4:** Genome assembly size by country. Boxplots represent distribution of genome assembly size for 38 distinct countries for which more than 1000 genome assemblies of SARS-CoV-2 are available in the GISAID database as of March 18th 2021. Countries are indicated on the y axis. Size of assembled genomes is reported on the x axis.

**Figure S5**: Comparison of several features for Pango lineages (in Figure S5A) and HGs (in Figure S5B). Boxplots display global distribution of each distinct feature described in Supplementary Table S12 and used to cluster SARS-CoV-2 Pango lineages and HGs. Blue: lineages/HGs associated with VOCs/VOIs. Red: remaining lineages/HGs. The acronym used to indicate every feature is reported in the title along with a Wilcoxon sum and rank test p-value measuring the level of significance of the difference between the 2 boxplots.

**Figure S6:** Distribution of genetic distance, computed as the total number of distinct polymorphic sites between closest pairs of genomes, assigned to Pango lineages/HGs associated to VOCs/VOIss and to all remaining Pango lineages/HGs (indicated as(“other”). Distributions are represented in the form of a violin plot. Distances are indicated on the Y axis.

## Supplementary Tables

**Supplementary Table S1**: List of the 93.265 allele variants identified in SARS-CoV-2 genomic sequences together with their functional annotation. POS: nucleotide position on the reference genome. REF: reference allele. ALT: alternative allele. annot: functional annotation according to CorGAT. HF_cumulative: does the variants reach an AF >= 1% when allele frequencies are computed cumulatively on all the genomic sequences: 1= TRUE. 0= FALSE. HF_regional: equivalent to HF_cumulative but with AF data computed at regional level. For variants associated with multiple regions, the maximum value is considered.

**Supplementary Table S2:** Genomes per country and correspondence with geographic macro-areas. Countries are reported in the first column. Geographic areas are indicated in the second column. See Figure 1 legend for the acronyms. The last 3 columns indicate the total number of reported cases, the incidence of COVID-19 (number of cases per 100.000 population) and the total number of Pango lineages detected in each country.

**Supplementary Table S3**: FDR for regional bias. Alleles are reported on the rows in the following format: <genomic position>_<reference>**|**>alternative>. Geographic areas are indicated in the columns. Acronyms of geographic regions according to Table S2. The minimum FDR value is reported in the last column

**Supplementary Table S4**: High quality assemblies. Percentage of high quality genomes associated with different countries. Column1: % based on original criteria defined in Chiara et al 2021. Column2: % based on revised criteria used in the current study. Column 3: difference between 1 and 2.

**Supplementary Table S5**: Complete list of genomes. Complete list of genomes considered in these analyses, including metadata derived from the GISAID database and assignment of HGs (last column) .

**Supplementary Table S6**: High Frequency allele variants associated with the 144 distinct haplogroups corresponding to the B.1.1.7 lineage. Variants are reported in the first column using the same format as in Table S3 (<genomic position>_<reference>**|**>alternative>).

Phenetic patterns of absence/presence (1= presence. 0= absence) are reported in a separate column for every haplogroup. Numbers in parentheses indicate the number of genomes assigned to each HG. The column “in Tot HG” reports the total number of HGs to which each variant is associated.

**Supplementary Table S7**: Contingency Table (number of genomes by country) of the first 50 genomes of each country, defined by isolation date (as reported in GISAID in date March 18th 2021), associated with each of the 144 HGs corresponding with B.1.1.7

**Supplementary Table S8**: Percentage(%) of the total number of the 144 HGs corresponding with B.1.1.7 submitted from distinct countries. Only countries that account for the submission of at least 1% of the genomic sequences of one or more HGs are reported. Countries are reported on the rows. HGs on the columns. The first row reports the total number of HGs that considered specific to each country according to our criteria

**Supplementary Table S9:** Sites under selection, determined by Hyphy (Kosakovsky-Pond et al^23^). Residue: amino acid residue. Fel: under selection according to Fel. True/False. Meme: under selection according to meme. True/False. Type: type of selection positive/negative.

**Supplementary Table S10:** Total number of genomes and proportion of high quality (HQ) genomes associated with each haplogroup. HGs with % HQ genomes < 65% are reported in bold. The last column indicates the “node purity” in the phylogenetic tree (see methods).

**Supplementary Table S11:** High Frequency variants associated with the 3 distinct haplogroups corresponding to the B.1.351 Pango lineage Phenetic patterns of absence/presence (1= presence. 0= absence) are reported in a separate column for every haplogroup. Numbers in parentheses indicate the number of genomes assigned to each HG. The last column reports the total number of HGs to which each variant is associated.

**Supplementary Table S12:** High Frequency variants associated with the 6 distinct haplogroups corresponding to the P.1 Pango lineage Phenetic patterns of absence/presence (1= presence. 0= absence) are reported in a separate column for every haplogroup. Numbers in parentheses indicate the number of genomes assigned to each HG. The last column reports the total number of HGs to which each variant is associated.

**Supplementary Table S13:** High Frequency variants associated with the 8 distinct haplogroups corresponding to the B.1.617.2 Pango lineage.

Phenetic patterns of absence/presence (1= presence. 0= absence) are reported in a separate column for every haplogroup. Numbers in parentheses indicate the number of genomes assigned to each HG. The last column reports the total number of HGs to which each variant is associated.

**Supplementary Table S14:** list of features used in the classification of SARS-CoV-2 lineages/HGs. Acronyms are reported in the first column. A brief description is reported in the second column

**Supplementary Table S15:** list of allele variants enriched in Pango or HG clusters associated with VOCs/VOIs. A FDR of 0.05 or below is used as the threshold for significant over-representation. C2lin: FDR for over-representation in Cluster 2, i.e., the cluster associated with VOCs and VOIs in the Pango lineages. C3HG: FDR for over-representation in Cluster 3 of HG, i.e., the cluster associated with VOCs and VOIs in the HG system. Column “Common”: 1= over represented in both C2lin and C3HG; 0= over represented only in one cluster (C2lin or C3HG). Annotation: functional annotation according to CorGAT

**Extended data** are available from the following Figshare repository: https://doi.org/10.6084/m9.figshare.14823366 . Please refer to the readme file for more informaion

